# The visual system of the genetically tractable crustacean *Parhyale hawaiensis*: diversification of eyes and visual circuits associated with low-resolution vision

**DOI:** 10.1101/527564

**Authors:** Ana Patricia Ramos, Ola Gustafsson, Nicolas Labert, Iris Salecker, Dan-Eric Nilsson, Michalis Averof

## Abstract

**Background:** Arthropod eyes have diversified during evolution to serve multiple needs, such as finding mates, hunting prey, and navigating in complex surroundings under varying light conditions. This diversity is reflected in the optical apparatus, photoreceptors and neural circuits that underpin vision. While this diversity has been extensively documented, our ability to genetically manipulate the visual system to investigate its function is largely limited to a single species, the fruitfly *Drosophila melanogaster*. Here, we describe the visual system of *Parhyale hawaiensis*, an amphipod crustacean for which we have established tailored genetic tools.

**Results:** Adult *Parhyale* have apposition-type compound eyes made up of ∼50 ommatidia. Each ommatidium contains four photoreceptor cells with large rhabdomeres (R1-4), expected to be sensitive to the polarisation of light, and one photoreceptor cell with a smaller rhabdomere (R5). The two types of photoreceptors express different opsins, belonging to families with distinct wavelength sensitivities. Using the *cis*.-regulatory regions of opsin genes, we established transgenic reporters expressed in each photoreceptor cell type. Based on these reporters, we show that R1-4 and R5 photoreceptors extend axons to the first optic lobe neuropil, revealing striking differences compared with the photoreceptor projections found in related crustaceans and insects. Investigating visual function, we show that *Parhyale* has a positive phototactic response and is capable of adapting its eyes to different levels of light intensity.

**Conclusions:** We propose that the visual system of *Parhyale* serves low-resolution visual tasks, such as orientation and navigation, based on broad gradients of light intensity and polarisation. Optic lobe structure and photoreceptor projections point to significant divergence from the conserved visual circuits found in other malacostracan crustaceans and insects, which could be associated with a shift to low-resolution vision. Our study provides the foundation for research in the visual system of this genetically tractable species.

## Introduction

Arthropods have elaborate visual systems that are capable of serving a wide range of tasks, including orientation and navigation, prey capture, habitat selection and communication, in diverse environments (see [1]). Depending on the life habits of individual species, the needs for spatial resolution, motion detection, sensitivity to colour or light polarisation, or vision in dim light will differ. This functional diversity is reflected in the anatomy of the eye and in the underlying neural circuits.

Compound eyes, the most common and best studied type of arthropod eye, are made up of multiple repeated units. Each unit, the ommatidium, consists of a light-focusing apparatus (including the cornea and crystalline cone cells) overlying a cluster of photoreceptor cells with a light-sensing rhabdom. Optical designs vary, encompassing eyes in which the ommatidia are optically isolated from each other (apposition type) or make up a common light-focusing unit (superposition type), where light is focused by reflection and/or refraction, and designs in which different areas of the eye have been specialised to perform different functions (reviewed in [2, 3]). Differences in the number of ommatidia - ranging from 1 to 30,000 – constrain the image resolution that can be achieved by a given eye, since each rhabdom corresponds to an image pixel [4], Within each ommatidium, differences in the types, morphology and arrangement of photoreceptors influence the capacity to detect colour and polarised light [1, 5].

At the molecular level, functional diversification is most evident in visual opsins, which largely determine the spectral sensitivity of each photoreceptor. Molecular phylogenetic analyses suggest that ancestral pancrustaceans had several distinct visual opsins belonging to the r-opsin family [6]. Extant crustaceans and insects have varying numbers of opsin genes, ranging from only one opsin in deep sea crustaceans with monochrome vision to >10 opsins with distinct spectral properties in stomatopod crustaceans and dragonflies [7-10].

Visual stimuli are processed in the underlying optic lobes. In malacostracan crustaceans and insects we can typically identify four distinct optic neuropils, named lamina, medulla, lobula and lobula plate [11-15]. A chiasm (crossing over of axons) is typically found between the lamina and medulla and between the medulla and the lobula [11, 13, 16].

Photoreceptors of each ommatidium connect to the optic lobe via short axons that terminate in the lamina (e.g. photoreceptors R1-6 in flies, R1-7 in decapods, named ‘short fibre’ photoreceptors) or extend longer axons that cross the lamina and terminate in the medulla (e.g. R7-8 in flies, R8 in decapods, named ‘long fibre’ photoreceptors) [17-19]. Diverse sets of local interneurons and projection neurons receive input from these two classes of photoreceptors and relay processed information to deeper neuropils of the optic lobe. In each neuropil, dedicated subcircuits perform distinct visual processing tasks, such as contrast enhancement, the detection of colour, polarised light or motion [15, 20-22].

Understanding the functional diversification of these neural circuits is challenging. Changes in the number of visual neuropils, their structure and their connections to each other have been documented in some groups (for example, branchiopod crustaceans have only two optic neuropils [11-13, 16, 23]), but the functional impact of these changes is not well understood. Moreover, functional changes in vision could be associated with subtle changes in neuronal subtypes and connectivity, which would be difficult to identify. Detailed studies of visual circuits have only been possible in a small number of experimental models. Most notable is the fruitfly *Drosophila melanogaster*, where neuroanatomical studies have been combined with powerful genetic tools and behavioural studies to yield a detailed understanding of some visual neural circuit architecture and function (e.g. [9, 20, 24-28]).

The array of genetic tools and resources currently available in *Drosophila* are unlikely to be matched in other species. However, some genetic tools and resources can undoubtedly be adapted to explore the functional diversification of the visual system in other species. For example, transgenic markers can serve to identify and to characterise specific cell types. Transgenes expressed in specific neuronal cells can be used to monitor and manipulate the activity of these cells or to ablate them, and in this way to investigate their roles within the neural circuits that underpin vision. A thorough understanding of the evolution of visual circuits requires the establishment of such tools in diverse species beyond insects.

Here, we focus on the visual system of *Parhyale hawaiensis*, an amphipod crustacean living in shallow marine habitats in the tropics (Figure 1A). Amphipods are widely distributed aquatic and semi-terrestrial crustaceans, belonging to one of the major evolutionary branches of crustaceans, the malacostracans. The visual system of amphipods remains relatively unexplored [29-33], though some species exhibit extreme adaptations (e.g. [34]). In recent years, a range of genetic tools and resources have been established in *Parhyale hawaiensis*, including transgenesis [35], CRISPR-mediated gene editing [36] and a sequenced genome [37], We exploit these resources to generate new tools for probing the anatomy and neural connectivity of the visual system of *Parhyale*. Our work reveals striking evolutionary diversification in visual circuits and provides the foundation for a genetics-driven analysis of visual function in this species.

**Figure 1.**
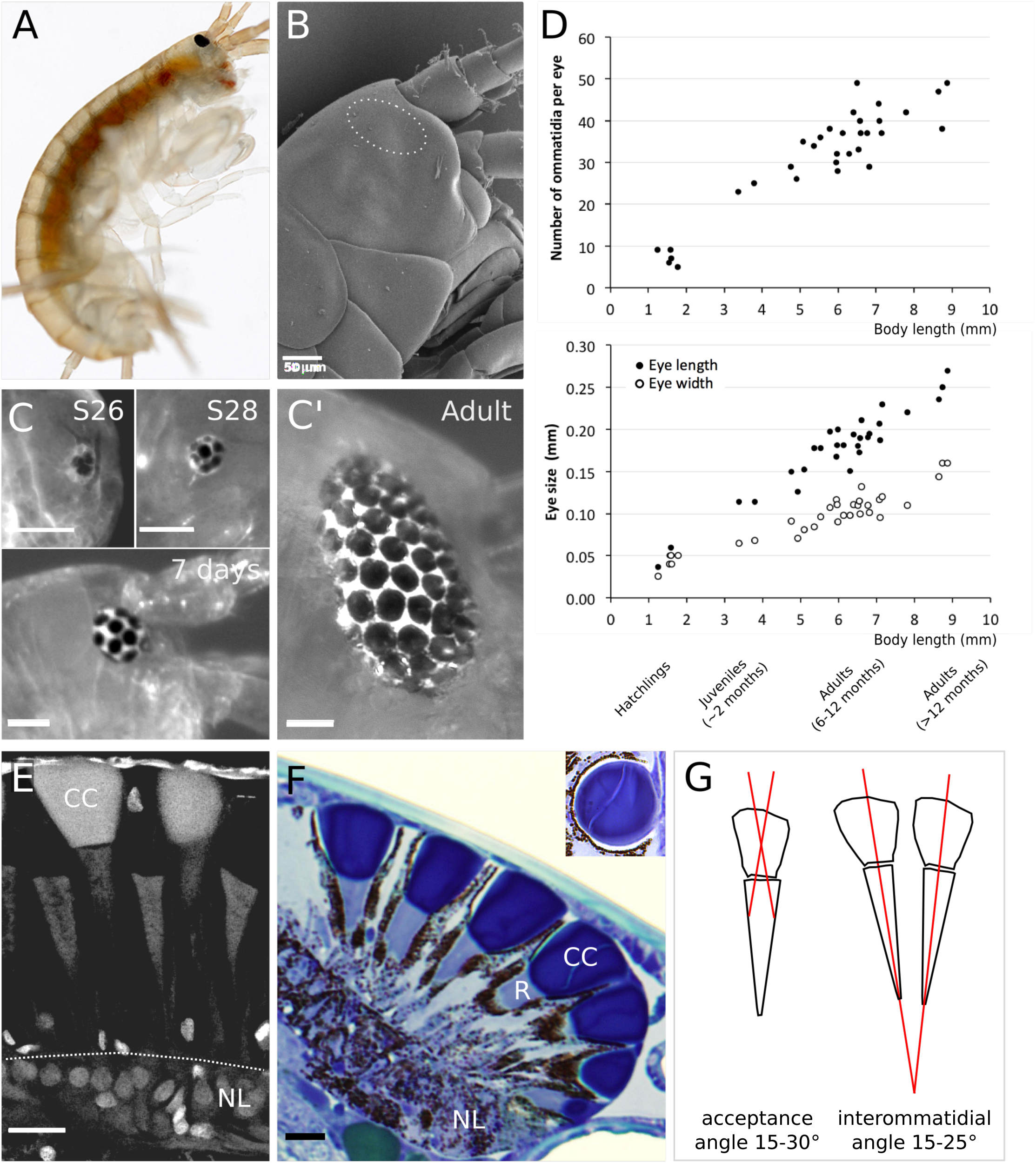
Growth and morphology of the compound eyes of *Parhyale*. **A**. Adult *Parhyale* with a dark pigmented compound eye typical of gammarid-like amphipods (photo credit: Vincent Moncorgé). **B**. Scanning electron micrograph of the head of a *Parhyale* hatchling. The surface of the cuticle is smooth, with no distinctive structure in the region covering the eye (marked by a dashed line). **C. C’.** Growth of the compound eye recorded in stage 26 and stage 28 embryos, a 7-day old juvenile and an adult. Scale bars, 50 μm. **D**. Growth of the compound eye during lifetime of *Parhyale*. The number of ommatidia per eye (top) and the length and the width of the eye (bottom) scale linearly with body length. **E**. Fluorescence image of a *Parhyale* compound eye, sectioned and stained with DAPI to reveal nuclei. The photoreceptor nuclei are concentrated at the base of the eye, below the rhabdoms. Crystalline cones show strong autofluorescence. Scale bar, 20 µm. **F**. Longitudinal semi-thin section through a *Parhyale* compound eye, stained with toluidine blue. The crystalline cone of each ommatidium (stained dark blue, CC) and the underlying rhabdom (stained light blue, R) are clearly visible, surrounded by dark pigment granules. The nuclei of the photoreceptor cells are located below the rhabdoms (NL). The inset photo shows a section through the crystalline cone, which consists of two cells. Scale bar, 20 µm. **G**. Illustration of acceptance and interommatidial angles measured from sections of adult *Parhyale* eyes. The acceptance angles of individual ommatidia are assessed from an optical nodal point in the middle of the crystalline cone and an assumed image plane located 1/3 of the distance down the rhabdoms (see Materials and Methods).

## Results

### Morphology and growth of the apposition-type compound eyes of *Parhyale*

*Parhyale* adults have a pair of dark-coloured oval or kidney shaped compound eyes located laterally on the head (Figure 1A, C). The ommatidia are arranged in rows, following the hexagonal packing typical of many compound eyes. The eyes first become visible in stage 25 embryos (staging according to [38]), typically consisting of 3 unpigmented ommatidia. These become pigmented and surrounded by additional ommatidia between stages 26 to 28 (Figure 1C). *Parhyale* hatch with approximately 8-9 mature (pigmented) ommatidia per eye. Further ommatidia are added gradually during their lifetime, accompanying the growth of the body, reaching approximately 50 ommatidia per eye in the oldest adults (Figure 1C’). The growth of the eye is anisotropic, starting from a round shape that gradually becomes elongated.

To quantify the growth of the eyes we measured eye size and ommatidial numbers in a sample of 30 juveniles and adults (Figure 1D). The surface area of the eye increases approximately 40-fold from hatchlings to adults (eye length increases from 0.037 to 0.270 mm, eye width from 0.026 to 0.160 mm), whereas the number of ommatidia increases approximately 6-fold in the same animals (from 8 to 49 ommatidia per eye). Hence, the surface occupied by each ommatidium increases approximately 7-fold from hatchlings to adults. These results suggest that *Parhyale* eyes keep growing due to a parallel increase in ommatidial number and ommatidial size.

To assess the internal anatomy of *Parhyale* compound eyes we produced semi-thin (2 µm) histological sections of adult heads embedded in epoxy resin. Longitudinal sections through the eye reveal the structure of ommatidial units, with a crystalline cone and an underlying rhabdom surrounded by dark granules (Figures 1F). Beneath the rhabdoms lies a layer containing a large number of nuclei, which likely include the nuclei of the photoreceptor cells (Figure 1E) [29]. The separation of each ommatidium by pigment granules and the direct contact of the crystalline cones with the underlying rhabdoms indicate that *Parhyale* eyes function as apposition-type compound eyes.

Semi-thin sections and scanning electron microscopy show that the cuticular cornea covering the eye is smooth and not divided into facets as in many other compound eyes (Figures 1B, F). The nearly flat and uniform cornea with constant thickness suggests that the cuticle does not play a role in focusing light and that *Parhyale* eyes are likely to perform equally well in aquatic and terrestrial environments (see [3]). Cross sections through the ommatidia reveal that the crystalline cone is formed by two cells (Figure 1F, inset), as is typically the case in amphipods [29].

As can be judged from the divergence of ommatidial axes in the semi-thin sections, the field of view of each adult compound eye is approximately 120° vertically and 150° horizontally. The interommatidial angles of adult eyes range between about 15° and 25°, and the acceptance angles of each ommatidium range from 15° to 30° (Figure 1G, see Materials and Methods). Acceptance angles define the ultimate resolution limit of a visual system, and they are expected to be similar to the sampling interval (interommatidial angles) if each ommatidium corresponds to a resolved visual pixel [3]. We thus infer that the number of ommatidia is also the number of resolved pixels the eye can use when there is sufficient light. These measurements suggest that *Parhyale* have a wide visual field but a notably low spatial resolution.

### Structure of ommatidia and arrangement of photoreceptors

Many insects and crustaceans possess 8 photoreceptors per ommatidium, but the precise number and the spatial arrangement of photoreceptors vary among different groups (reviewed in [5, 39, 40]). To assess the number, morphology and organisation of photoreceptors within the ommatidia of *Parhyale*, we produced ultra-thin sections of adult heads and examined these by transmission electron microscopy (TEM).

Longitudinal and transverse sections through the ommatidia revealed that each rhabdom is surrounded by electron-dense vesicles, corresponding to pigment granules within the photoreceptor cells (Figure 2). Furthermore, transverse sections show that each ommatidium is composed of five photoreceptor cells: four with large rhabdomeres (named R1-4) and one photoreceptor with a smaller rhabdomere (named R5) (Figure 2A). The rhabdomeres of all the photoreceptors are closely apposed to each other, forming a closed or fused rhabdom [41, 42], Unlike in some other crustaceans with fused rhabdoms (e.g. [43]), in *Parhyale* the microvilli of different rhabdomeres do not interdigitate with each other, but occupy distinct sections of the rhabdom throughout its length (Figure 2B, C). The rhabdomeres of all photoreceptors contribute to the entire length of the rhabdom (Figure 2D, E).

**Figure 2.**
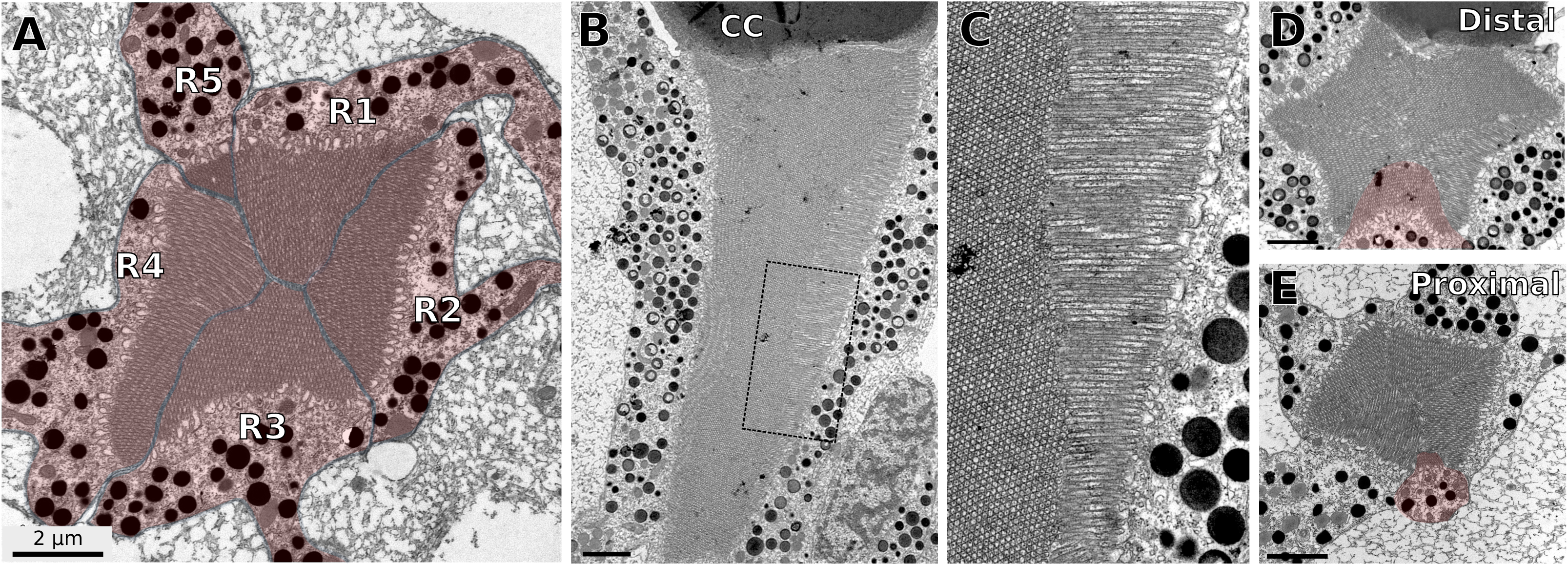
Photoreceptor arrangement in the ommatidia of *Parhyale*. **A**. Transmission electron micrograph of a transverse section through a *Parhyale* ommatidium at the level of the rhabdom. The rhabdom, at the centre of the image, consists of the tightly packed microvilli of photoreceptor cells. Five photoreceptor cells contribute to the rhabdom, numbered clockwise R1-5 (highlighted in red). Dark granules are visible in the cytoplasm of each photoreceptor cell. The photoreceptors are surrounded by lighter stained material, which belongs to accessory cells with reflective granules. **B**. Transmission electron micrograph of a longitudinal section through a *Parhyale* ommatidium. The rhabdom is visible below the crystalline cone (labelled CC) and surrounded by the cytoplasm of photoreceptor cells containing dark granules. The labelled square corresponds to an area shown at higher magnification in panel C. **C**. Higher magnification view of the longitudinal section shown in panel B. The microvilli of neighbouring photoreceptors are arranged perpendicular to each other, and retain the same orientation throughout the length of the rhabdom. **D. E.** Transmission electron micrographs of transverse sections at two different levels of the rhabdom: at the distal end of the rhabdom, close to the crystalline cone **(D)**, and at a proximal level near the nuclear layer **(E).** The R5 photoreceptor (highlighted in red) contributes to the rhabdom at both levels. Scale bars, 2 µm.

Longitudinal sections show that the microvilli of a given photoreceptor remain well aligned with each other throughout the length of the rhabdom (Figure 2B, C). This arrangement suggests that each photoreceptor is sensitive to a particular direction of light polarisation [44-46]. The microvilli of R1 and R3 are oriented parallel to each other and orthogonally to those of R2 and R4 (see Figure 2A, C), suggesting that, within each ommatidium, photoreceptors R1+R3 and R2+R4 have differential sensitivities with respect to the direction of light polarisation.

### Photoreceptors R1-4 and R5 express different opsins

Opsins determine the spectral sensitivity of photoreceptors. To identify which opsins are expressed in *Parhyale* compound eyes we searched for opsin homologues in *Parhyale* embryo and adult head transcriptomes [37, 47, 48] (see Materials and Methods). We identified two opsins, named *PhOpsin1 and PhOpsin2. In situ* hybridization confirmed that both are expressed in the eyes of *Parhyale* from stage 26 embryos to adults. *PhOpsin1*. is expressed widely in the retina, whereas *PhOpsin2* is expressed in a punctate pattern localised to one side of each rhabdom (Figure 3 insets).

**Figure 3.**
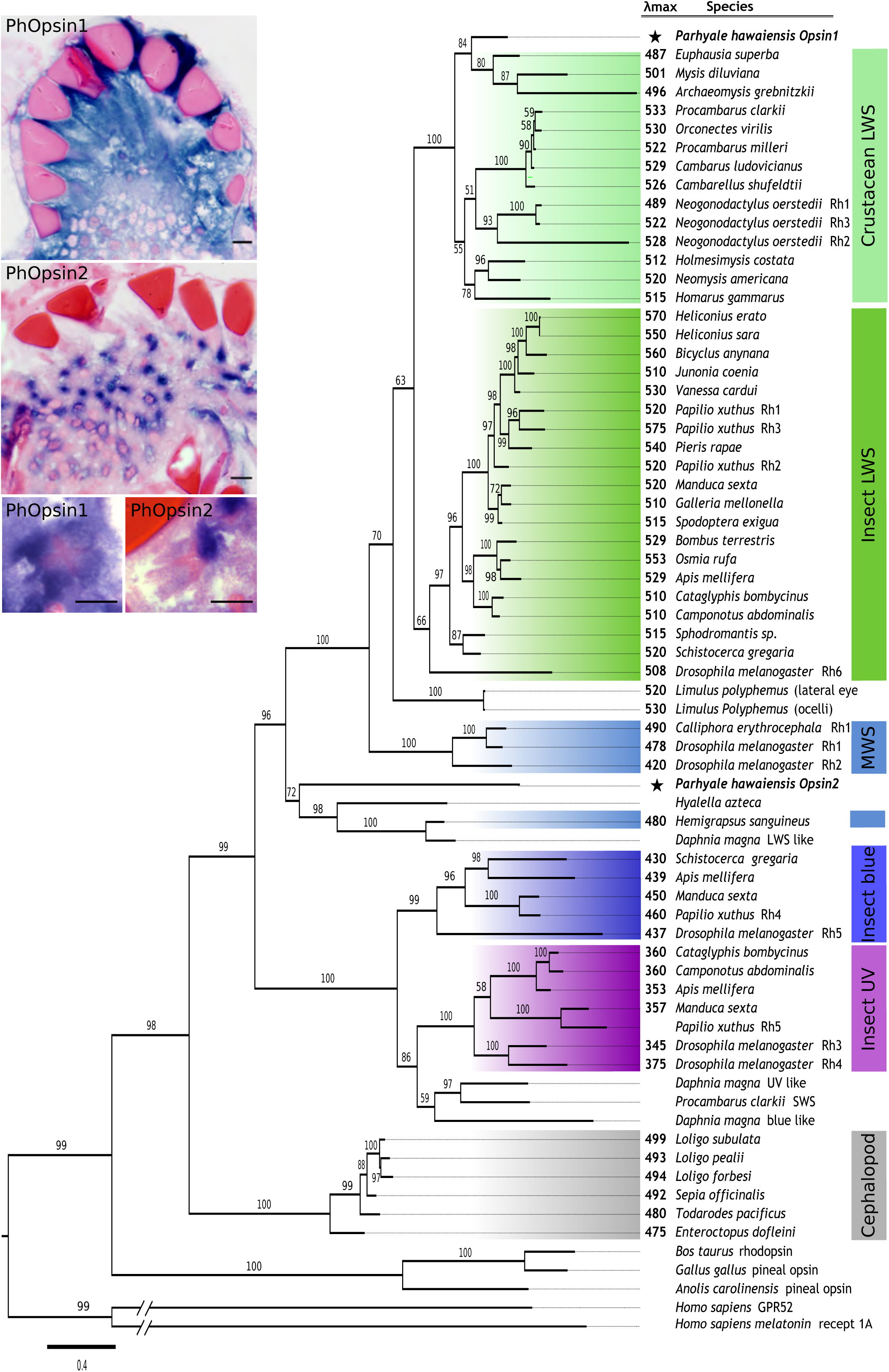
Phylogenetic relationships and expression pattern of *Parhyale* opsins Maximum-Likelihood tree depicting the relationships of *PhOpsin1* and *PhOpsin2* with the major families of pancrustacean r-opsins (see Material and Methods). Insets show the expression of *PhOpsin1* and *PhOpsin2* mRNA by *in situ* hybridization in the eyes of adult *Parhyale* (purple-blue colour). *PhOpsin1* is expressed widely in the retina, whereas *PhOpsin2* is expressed in a punctate pattern. High magnification views of individual rhabdoms (lower panels, rhabdoms stained pink) show that *PhOpsin1* expression is widespread whereas *PhOpsin2* expression is localised to one corner of the rhabdom. Tissue sections were stained with basic fuchsin and eosin-erythrosine. Scale bars, 10 µm.

Phylogenetic analysis of arthropod opsin sequences shows that *PhOpsin1* is most closely related to insect and crustacean long-wavelength-sensitive (LWS) opsins, which have absorbance maxima at 490-530 nm, whereas *PhOpsin2* is most closely related to a crustacean middle-wavelength-sensitive (MWS) opsin, which has an absorbance maximum of 480 nm (Figure 3, [49]). These relationships suggest that PhOpsin1 and PhOpsin2 are likely to confer distinct spectral sensitivities to the photoreceptors in which they are expressed.

To determine the photoreceptors in which each opsin is expressed and to generate markers for each photoreceptor type, we set out to identify *cis*.-regulatory elements that drive the expression of each opsin. For this we generated reporter constructs by cloning genomic regions upstream of the start codon of each opsin gene and placing them upstream of a membrane tagged fluorescent marker gene (see Materials and Methods). These reporter constructs were inserted in the *Parhyale* genome by *Minos*.-mediated transgenesis [35].

From *PhOpsin1* we cloned a 1.6 kb genomic fragment that includes the 5’ UTR, promoter region and upstream sequences (Figure 4A), and placed this fragment upstream of the coding sequence of membrane-tethered EGFP-CAAX. This *PhOpsin1:EGFP-CAAX*. reporter gave a strong fluorescent signal in the eyes of *Parhyale* from stage 27 embryos to adults. For *PhOpsin2*, we found that sequences up to 5 kb upstream from the start codon were insufficient to drive fluorescent protein expression, but a 3.1 kb genomic fragment that also includes the first intron of *PhOpsin2* (Figure 4A) gave robust expression of mKate-CAAX in the eyes. This *PhOpsin2:mKate-CAAX* reporter was expressed in the eyes from stage 28 embryos to adults.

**Figure 4.**
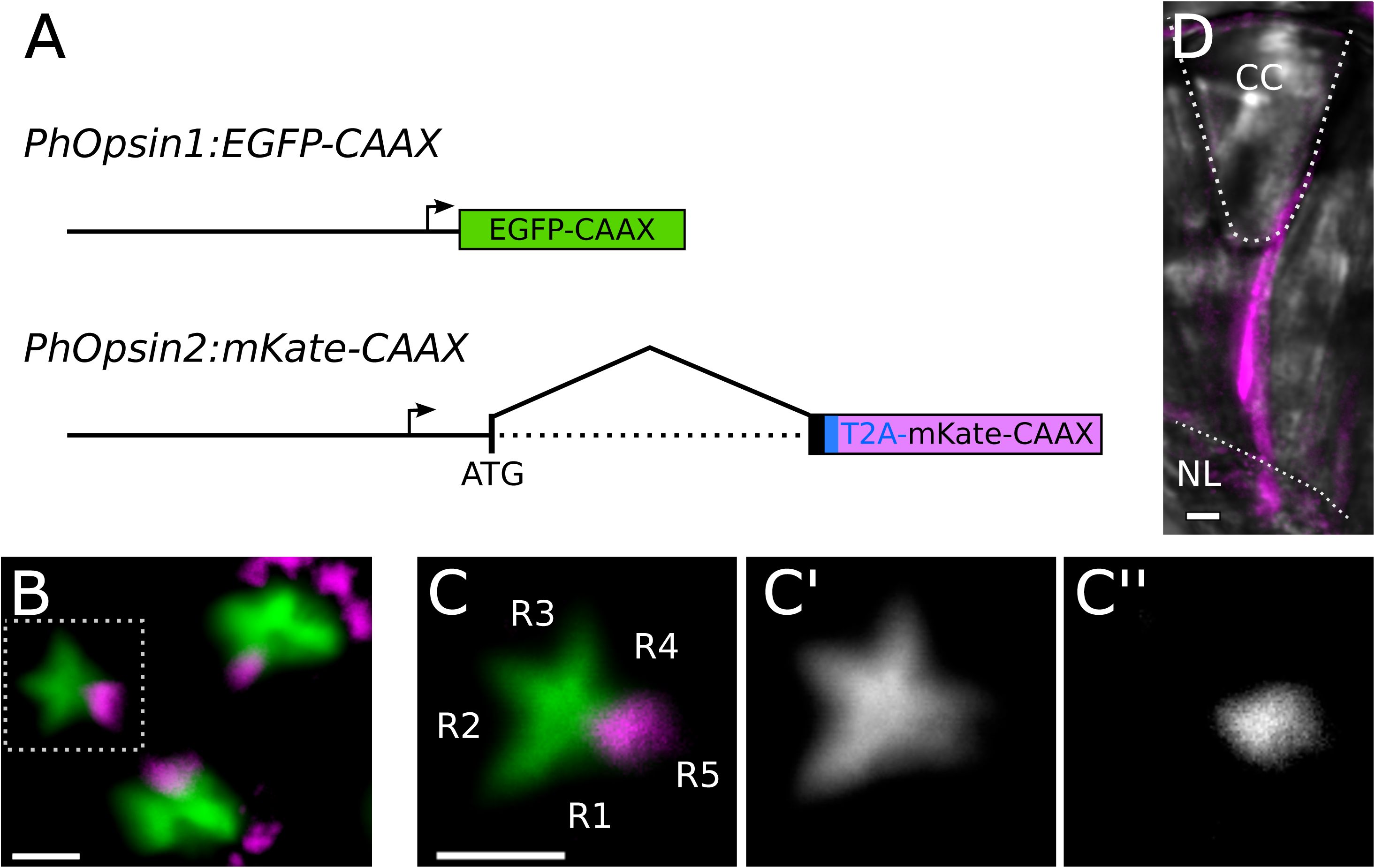
*Parhyale* opsin reporters. **A**. Design of the *PhOpsin1:EGFP-CAAX* and *PhOpsin2:mKate-CAAX* reporters. The genomic fragments of *PhOpsin1* and *PhOpsin2* are 1.6 and 3.1 kb long, respectively, including upstream sequences, promoters (arrow) and 5’ UTRs. The *PhOpsin2:mKate-CAAX* reporter includes the endogenous start codon of *PhOpsin2* (ATG), part of the coding sequence of *PhOpsin2* (interrupted by an intron) and the T2A self-cleaving peptide sequence (in blue), both cloned in frame with the mKate-CAAX coding sequence. **B**. Fluorescence microscopy in a live transgenic embryo carrying both reporters. Three ommatidia are visible in the field of view, with EGFP-CAAX (green) and mKate-CAAX (magenta) fluorescence localised in the rhabdom. The mKate channel also shows autofluorescence of pigment granules. **C-C”.** Higher magnification view of a single ommatidium, revealing the star-shaped pattern of the rhabdom. EGFP-CAAX (green) localises in most of the rhabdom, whereas mKate-CAAX (magenta) is restricted to one tip. **D**. Longitudinal view of an ommatidium of a transgenic adult carrying the *PhOpsin2:mKate-CAAX* reporter. The structure of the ommatidium is revealed by brightfield microscopy (in grey). The mKate signal (magenta) extends through the entire length of the rhabdom, from proximal (near the nuclear layer, NL) to distal (near the crystalline cone, CC). Scale bars, 5 µm.

Live microscopy of the eyes in late embryos revealed that CAAX-tagged EGFP and mKate are predominantly localised in the rhabdoms of each ommatidium that consist of densely stacked cell membranes (Figure 4B, C). The two reporters are expressed in distinct patterns, which are consistent with the results on opsin expression described earlier: *PhOpsin1*-driven EGFP-CAAX is localised in a star-shaped pattern which resembles the fused rhabdomeres of photoreceptors R1-4 (Figure 4C’), whereas *PhOpsin2*-driven mKate-CAAX is localised in a spot at one end of the star-shaped pattern, which likely corresponds to the rhabdomere of photoreceptor R5 (Figure 4C”). In sections of adult eyes, mKate-CAAX extends from the crystalline cone to the nuclear layer (Figure 4D), indicating that the smaller rhabdomere of photoreceptor R5 spans the entire length of the rhabdom.

These observations indicate that photoreceptors R1-4 and R5 have distinct colour sensitivities, determined by the spectral properties of PhOpsin1 and PhOpsin2, respectively.

### Structure of the optic lobes of *Parhyale*

Visual signals generated in the retina are processed in the underlying optic lobe, which in arthropods is composed of distinct neuropils. To examine the structure of the *Parhyale* optic lobe, we prepared toluidine-blue stained semi-thin sections of adult brains. We also performed immunostainings with antibodies for acetylated tubulin and synapsin to visualise axons and synapse-rich neuropils, respectively, in embryonic and adult brains (Figure 5).

**Figure 5.**
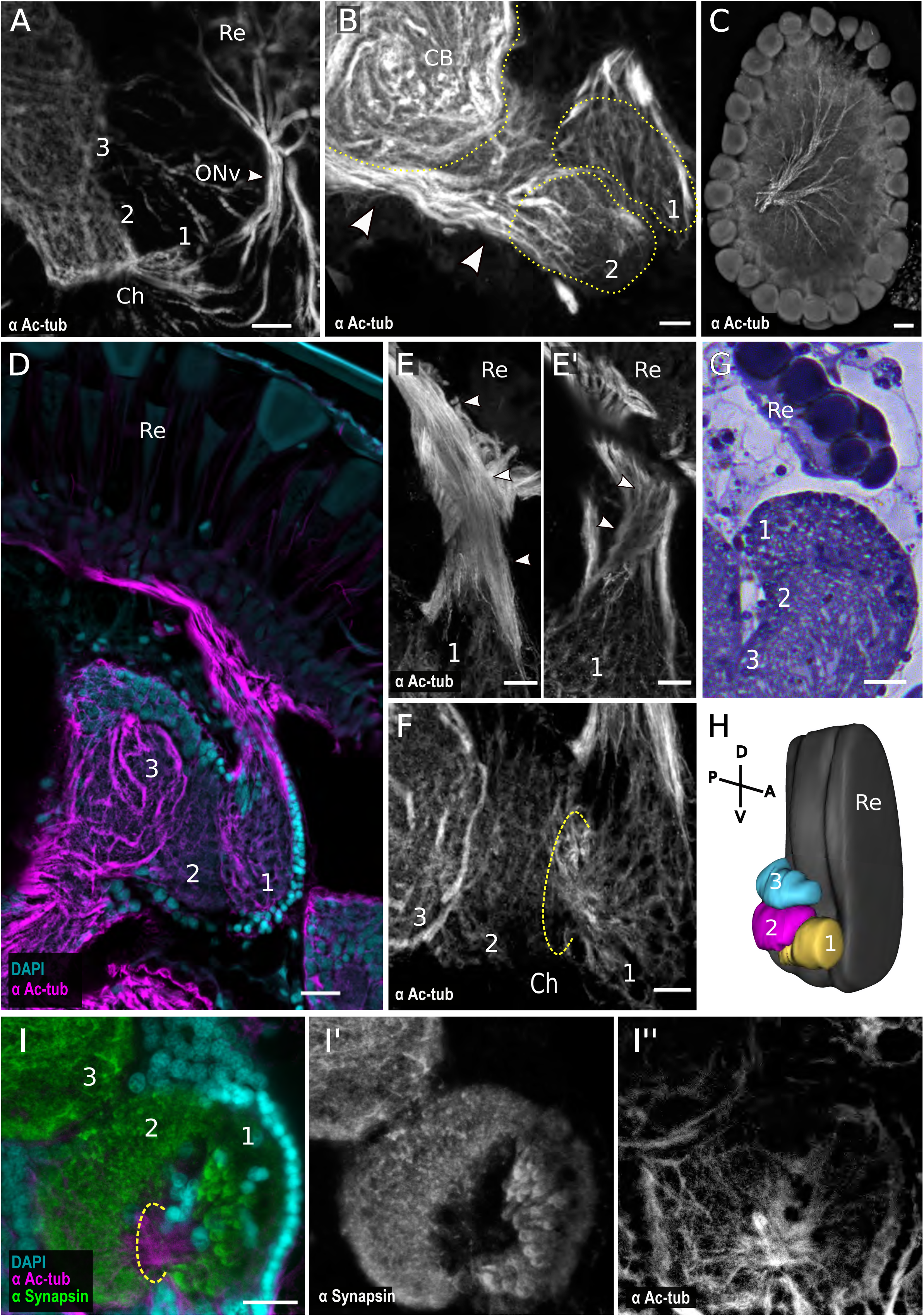
Architecture of *Parhyale* optic lobes. **A**. Whole mount stage 27 embryonic brain stained with an antibody for acetylated tubulin. The optic nerve (ONv) connects the retina (Re) to the first optic neuropil (1). A chiasm (Ch) is present between the first (1) and the second (2) neuropils. A third neuropil (3) is also distinguishable. Scale bar, 10 µm. **B**. Section of an adult brain stained with acetylated tubulin antibody. Higher order neurons (arrowheads) connect the second optic neuropil (2) and the central brain (CB). Scale bar, 10 µm. **C**. Adult retina stained with acetylated tubulin. Photoreceptor axons converge at the back of the retina forming the optic nerve. Scale bar, 20 µm. **D**. Section of an adult brain stained with DAPI and with an antibody for acetylated tubulin. Photoreceptor axons emerge from the retina (Re) and converge ventrally forming the optic nerve which connects to the first neuropil (1). The second (2) and third (3) neuropils are also visible. Scale bar, 20 µm. **E-E’.** Detailed view of the optic nerve (same specimen as in panel D) imaged at different tissue depths, suggesting a potential chiasm. Axons emerging from the retina connect to the anterior/frontal part of the first optic neuropil (arrowheads in E). Focusing 10 µm deeper within the optic nerve reveals axons connecting to the posterior part of the first neuropil (arrowheads in E’). Scale bar, 10 µm. **F**. Detailed view of the connections between optic neuropils 1, 2 and 3 (same specimen as in panel D), forming a chiasm (Ch) between neuropils 1 and 2. Scale bar, 10 µm. **G**. Horizontal semi-thin section through a *Parhyale* adult brain, stained with toluidine blue, showing part of the retina (Re) and the three optic neuropils. Scale bar, 20 µm. **H**. 3D reconstruction of the retina and optic lobe of *Parhyale* based on whole mount adult brain stained with an antibody for acetylated tubulin. The first neuropil (1) is located anteriorly and ventrally relative to the other neuropils. **1-1”.** Whole mount adult *Parhyale* brain stained with DAPI and with antibodies for acetylated tubulin and Synapsin. Three distinct neuropils can be distinguished (labeled 1, 2 and 3). Synapsin staining is strongest in the inner part of the first neuropil (I’). The chiasm between the first and second neuropil is visible (I”). Scale bars, 20 µm.

Acetylated tubulin staining of the entire brain in stage 27 embryos reveals the axons projecting from the newly formed eyes to the optic lobes of the brain. These axons extend considerably to reach the first visible neuropil, forming an axon bundle that has been referred to as the optic nerve [31] (Figure 5A). A second and third neuropil are also distinguishable in these stainings. The first and second neuropils are connected to each other through a chiasm, but connections to the third optic neuropil do not seem to involve a crossover of axons (Figure 5A).

In immunostainings of thick (150-300 µm) vibratome sections through *Parhyale* adult brains, three distinct neuropils are seen in close proximity with each other (Figures 5D and 5H). As in the embryonic brain, the first and second neuropils (but not the second and the third) are connected through a chiasm (Figure 5F). Unlike in insects and other malacostracan crustaceans where the first optic neuropil (the lamina) is closely apposed to the retina, in *Parhyale* axons can be seen emerging from the back of the retina and converging to form an optic nerve which extends to the first neuropil (Figures 5C, D). These axons appear to cross each other, before reaching the first optic neuropil, forming a putative chiasm (Figure 5E, E’). Some neurons can be seen connecting the second neuropil with the central brain (Figure 5B). Toluidine-blue stained semi-thin sections confirm this overall structure of the optic lobes (Figure 5G).

In dissected adult brains, acetylated tubulin and synapsin stainings reveal a similar overall organisation of the optic lobe (Figure 51-1”). Synapsin stainings reveal that the first optic neuropil is cup-shaped, and its inner part is organised in repeated units that could reflect a columnar organisation (Figure 51’).

### All photoreceptors project to the first optic neuropil

Depending on their inputs to visual processing tasks, different photoreceptor cells may project their axons to different optic neuropils. Typically, in malacostracan crustaceans and insects, photoreceptors either make short projections that terminate in the lamina (e.g. R1-6 in flies, R1-7 in decapods and stomatopods) or long projections that cross the lamina and terminate in the medulla (e.g. R7-8 in flies, R8 in decapods and stomatopods) [17-19, 50].

We used the opsin reporters described earlier to visualise the axonal projections of *Parhyale* photoreceptors R1-4 (*PhOpsin1:EGFP-CAAX* reporter) and R5 *(PhOpsin2:mKate-CAAX* reporter) in stage 28 embryos and adults. Live imaging of the embryonic brain shows that photoreceptor projections labelled by both reporters coalesce into the optic nerve and extend to the first optic neuropil (Figure 6A). The axons from both types of photoreceptors terminate at the first optic neuropil. No axons were detected extending further.

**Figure 6.**
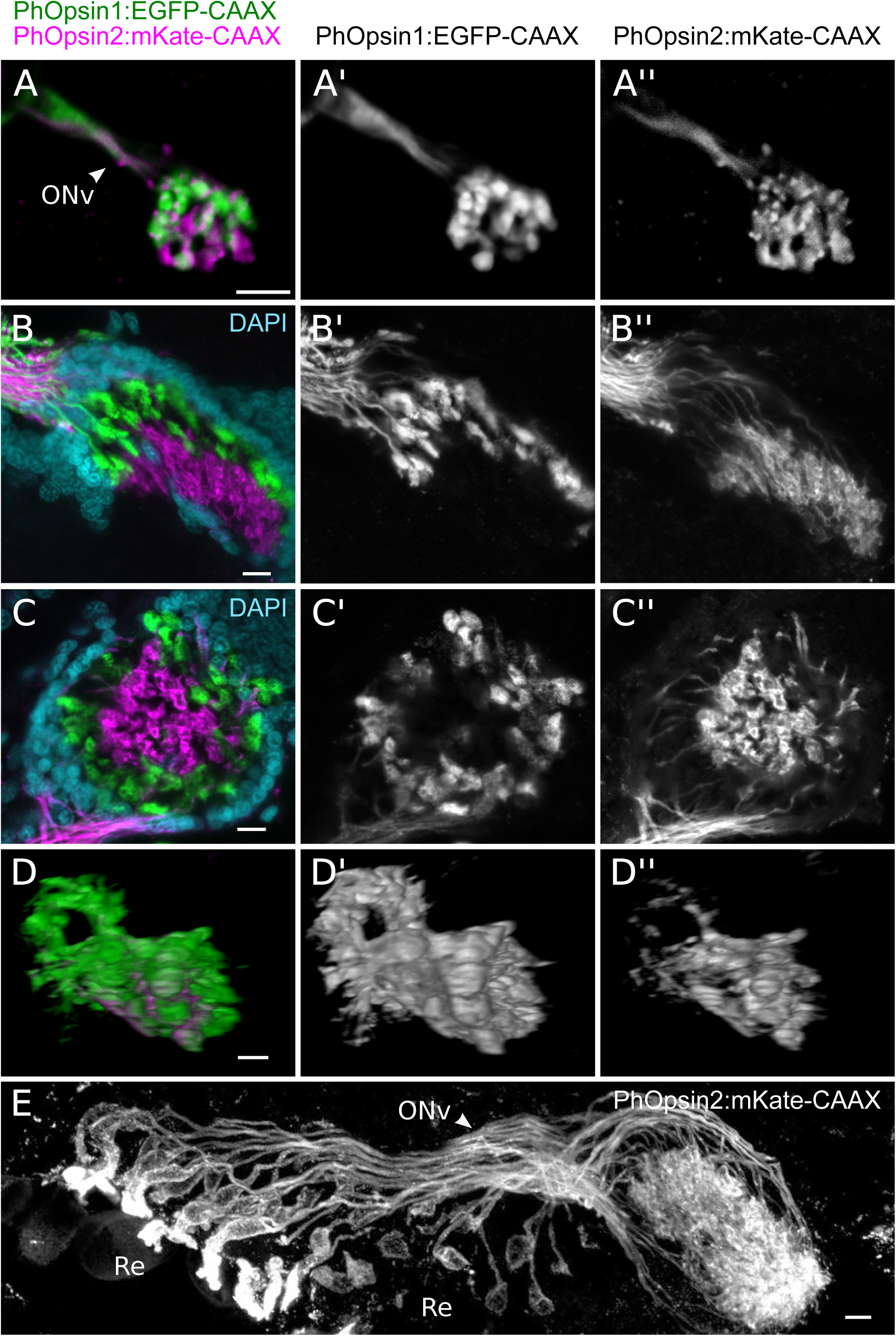
Photoreceptor projections visualised using opsin transgene reporters. **A**. Confocal imaging of a live transgenic embryo carrying the *PhOpsin1:EGFP-CAAX* and *PhOpsin2:mKate-CAAX* reporters. Photoreceptor axons labelled by EGFP-CAAX (green) or mKate-CAAX (magenta) form the optic nerve (ONv) and terminate in the same neuropil. **B**, **C.** Confocal imaging of the first neuropil of adult *Parhyale* carrying the *PhOpsin1:EGFP-CAAX* and *PhOpsin2:mKate-CAAX* reporters and stained with DAPI (in cyan). Photoreceptor projections labelled by EGFP-CAAX (green) terminate in the outer layer of the cup-shaped first neuropil, surrounding photoreceptor projections labelled by mKate-CAAX (magenta) which terminate at the interior. **D**. 3D rendering of a confocal scan through the first optic neuropil of the brain of an adult *Parhyale* carrying both opsin reporters. Photoreceptor axons labelled by EGFP-CAAX and mKate-CAAX project to the first optic neuropil and do not extend to the second neuropil (located on the right). The optic nerve is located at the left of the image. **E**. Lower magnification view of *PhOpsin2:mKate-CAAX* labelled photoreceptors, including the photoreceptor cell bodies at the retina (Re), the optic nerve (ONv) and the first optic neuropil (on the right of the image). Photoreceptor axons appear to cross each other before reaching the first neuropil. Scale bars, 10 µm.

To examine whether these projections are maintained in adults, the brains of double transgenic adults were fixed, dissected and stained with antibodies for acetylated tubulin, EGFP and mKate. Similarly to embryos, we observed that all photoreceptor projections terminate in the first optic neuropil (Figure 6B-D, 5 adults analysed). In stainings of adult brains it is clear that the *PhOpsin1-* and *PhOpsin2-* expressing photoreceptors terminate in different layers of the first optic neuropil (Figure 6B, C). The axons of *PhOpsin1*-expressing photoreceptors terminate in the outer layer of the cup-shaped neuropil. The axons of *PhOpsin2-expressing* photoreceptors cross that outer layer and terminate in the inner part of the neuropil. Some axons labelled by the *PhOpsin2:mKate* reporter appear to cross each other before reaching the first neuropil, confirming our observations of a potential chiasm described earlier (Figure 6E).

These results suggest that, unlike what has been reported in insects and other malacostracan crustaceans, the projections of all photoreceptors in the eyes of *Parhyale* terminate in the same optic neuropil.

### Adaptation to light intensity and phototactic responses

To start probing the physiology and function of *Parhyale’s* visual system, we have made some initial observations on two functional responses mediated by light: the adaptation of eyes to different intensities of ambient light and simple phototactic responses.

The compound eyes of arthropods typically adapt to different levels of light intensity by controlling the amount of light that reaches the rhabdoms. This adaptation can be achieved by redistribution of dark and reflective pigment granules within each ommatidium and/or by changes in the morphology and size of the rhabdom [51-53]; these changes are also influenced by endogenous circadian rhythms [54], In *Parhyale*, we have observed that dark-adapted eyes have a bright reflection surrounding the ommatidial facets, whereas light-adapted eyes are more darkly pigmented. The reflection fades away within a few minutes when the animals are transferred from a dark to a bright environment (Figure 7A), most likely reflecting shifts in the distributions of dark and reflective pigment granules within the eyes. This change can be observed equally during day and night.

**Figure 7.**
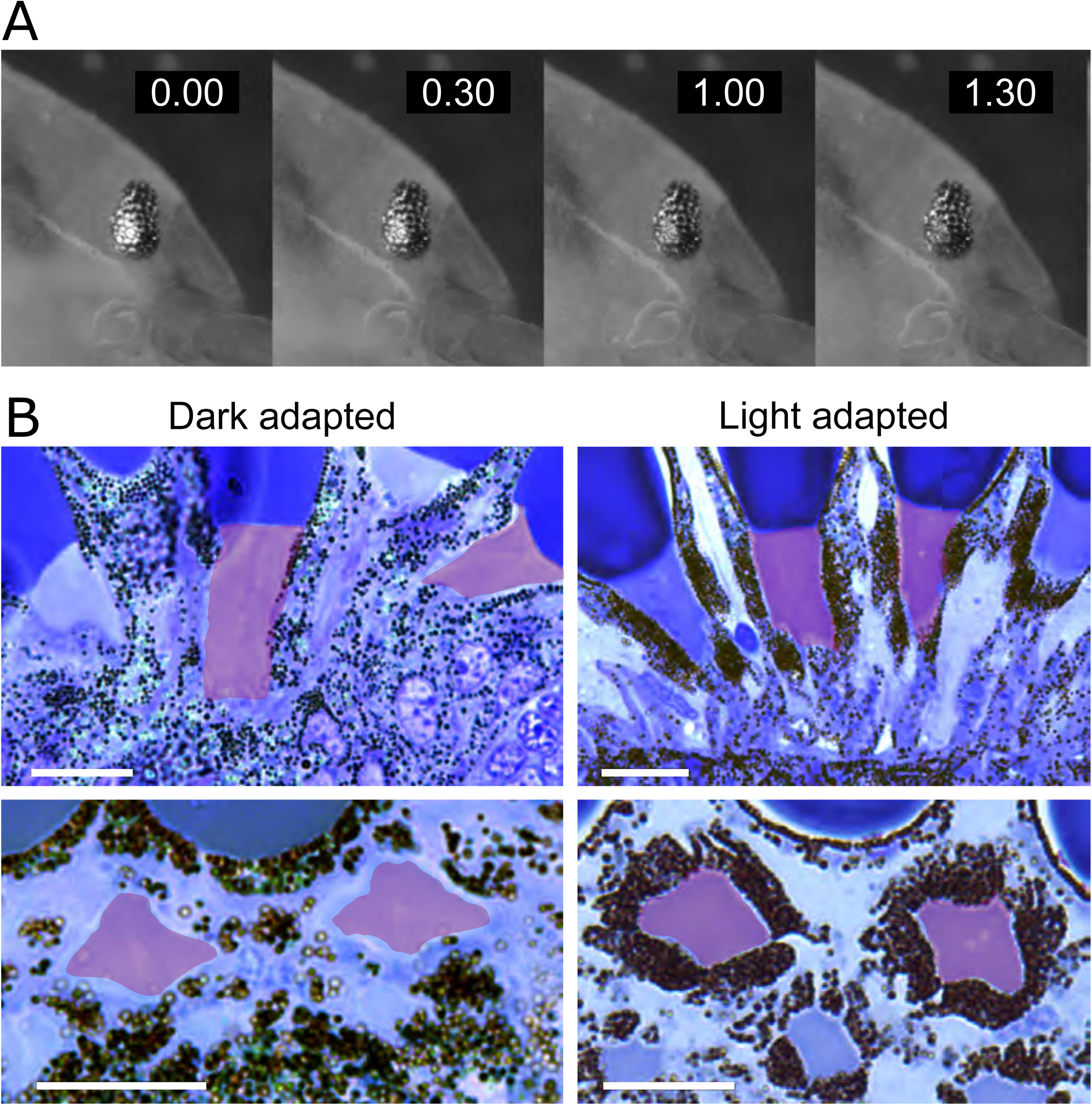
Adaptation of the eyes of *Parhyale* to light intensity. **A**. Dark-adapted eye of adult *Parhyale* imaged at different times after transfer to a bright environment. The reflection surrounding the ommatidial facets diminishes rapidly in response to light exposure. **B**. Semi-thin sections of dark-adapted and light-adapted eyes of adult *Parhyale*. Dark pigment granules are dispersed in dark adapted eyes, but concentrate around the rhabdom in light-adapted eyes (rhabdoms are highlighted in red). Longitudinal and transverse sections through ommatidia are shown in the top and bottom panels, respectively. Scale bars, 20 µm

To understand these changes in pigment distribution, we examined sections of dark- and light-adapted *Parhyale* eyes. In light-adapted animals, dark pigment granules are seen concentrated near the rhabdom, whereas in dark-adapted animals they disperse away from the rhabdom (Figure 7B; reflective granules are not visible in these preparations). The morphology and the size of the rhabdoms appear unchanged.

These results indicate that the eyes of *Parhyale* can adapt to different levels of ambient light by changing the distribution of dark pigment granules within the photoreceptor cells, to modulate the exposure of rhabdoms to the incoming light.

To explore whether *Parhyale* show phototaxis, we performed a simple two-choice behavioural assay. Adult *Parhyale* were placed at the centre of a T-maze with the choice of following a dark or an illuminated extremity of the maze. The experiment was conducted with different light intensities. Under low light intensities (<400 lux), 55% of the animals (182 out of 330) moved to the illuminated extremity of the maze, whereas 27% (89 out of 330) moved to the dark extremity (chi-squared p<0.001). In more intense light (>700 lux), 31% of animals (31 out of 100) moved to the illuminated extremity and 46% (46 out of 100) moved to the dark extremity (chi-squared p=0.088). These results indicate that *Parhyale* are positively phototactic at low light intensity, but not at high light intensity.

## Discussion/Conclusions

The optical apparatus of *Parhyale* follows the typical design of apposition-type compound eyes, where each ommatidial facet transmits light to just one underlying rhabdom. *Parhyale* eyes have a small number of ommatidia but sample a large field of view. Thus, in large adults, each eye contains approximately 50 ommatidia and samples a visual space that measures approximately 120° x 150°, with a sampling angle of 15-30° per ommatidium. This results in limited spatial resolution, the equivalent of taking a panoramic photo at 50-pixel resolution. This level of resolution is likely shared with other gammarid-like amphipods that have similar eyes. It comes in sharp contrast with the visual acuity of most other malacostracan crustaceans and insects, which is typically one to two orders of magnitude higher [4, 55-57].

Low resolution places constraints on the visual tasks that *Parhyale* can perform. For example, *Parhyale* are unlikely to be able to locate their mates or to find food using vision [58]; anecdotal evidence from gammarids supports this (e.g. [59]). However, low spatial resolution would be adequate for finding suitable habitats and for positioning within the habitat [58, 60]. Indeed, previous experiments on talitrid amphipods [32, 61] and the results of our phototaxis experiments suggest that *Parhyale* are able to perform such tasks.

The number and size of ommatidia increase in parallel with the growth of the body during the lifetime of *Parhyale*. As noted previously for isopod crustaceans [62], these changes are likely to influence both the visual acuity of the eyes (related to ommatidial numbers) and their sensitivity (related to ommatidial size). In the future, it will be interesting to investigate how new ommatidia are incorporated into the eyes, gradually extending the visual field and retinotopic maps in the context of a functioning visual system, and how the changes in visual performance impact visually-guided behaviours in juvenile and adult stages.

The structure of *Parhyale* eyes – including the number, arrangement and types of photoreceptors – is very similar to the pattern previously described in gammarid amphipods [29]. Thus, in spite of its low resolving power, this type of eye appears to have been conserved for at least 100 million years of divergent evolution (divergence estimate for gammarids and talitrids obtained from http://www.timetree.org/). Unlike what has been suggested in a previous study on talitrids [32], we find that the rhabdomeres of all photoreceptors (including R5) contribute to the entire length of the rhabdom.

The structure and arrangement of rhabdomeres within the eyes of *Parhyale* suggest that photoreceptor pairs R1+R3 and R2+R4 have differential sensitivities to light polarisation, and thus that visual perception in this species could be influenced by polarised light. Aquatic animals that are sensitive to light polarisation can use this information to enhance contrast by subtracting scattered light, enabling them to navigate or to orient themselves based on the celestial pattern of light polarisation [46, 63, 64], Outside of the water, animals can also make use of polarised reflected light to detect the surface of water [65]. While contrast enhancement is unlikely to be relevant for *Parhyale*, since their low resolution vision will not allow them to detect small- or medium-sized objects, orientation and navigation based on polarised light are likely to be relevant. As a member of the talitrid amphipods, which include the semi-terrestrial sandhoppers, sensitivity to polarised light may be relevant both inside the water (e.g. to detect the shore, [66]) and outside (to find water, [65]).

To detect light polarisation, the visual system of *Parhyale* would need to compare the signals received by photoreceptor pairs R1+R3 and R2+R4 of each ommatidium. In stomatopod crustaceans polarisation-sensitive photoreceptors project their axons to the lamina [50], whereas in insects they project to the medulla [5]. In *Parhyale*, R1+R3 and R2+R4 all project to the first optic neuropil. In the future it will be relevant to understand how their inputs are processed: whether they are compared (as would be required for polarised light detection) or added (cancelling polarisation sensitivity).

The presence of two opsins expressed in distinct sets of photoreceptors raises the possibility that *Parhyale* have dichromatic vision. Although the specific spectral properties of each opsin can only be determined by direct measurements of the wavelength of maximal absorbance, the finding that PhOpsin1 and PhOpsin2 are most closely related to opsins with long- and mid-wavelength sensitivities, respectively, suggests their most likely absorbance spectra. This is consistent with previous work on talitrid amphipods which revealed two peaks of sensitivity, at 520 and 430 nm [32, 33].

In insects and malacostracan crustaceans, the first step in processing colour information takes place in the medulla, which receives and compares inputs from photoreceptors with different spectral sensitivities. The input comes directly from long photoreceptor projections (from R7-8 in *Drosophila* and R8 in malacostracans) and indirectly from the other photoreceptors via higher-order neurons that relay their signals from the lamina to the medulla [9, 17-20, 50, 67], One of the most unexpected results of our study has been the discovery that in *Parhyale* all photoreceptor axons terminate in the first optic neuropil. To our knowledge this has not been observed in other malacostracan crustaceans or insects. However in some species, including more distant branchiopod crustaceans, photoreceptors of different spectral sensitivities do terminate in the first neuropil [50, 68, 69]. Colour discrimination in *Parhyale*, if it exists, would require the different spectral inputs from R1-4 and R5 to be compared directly in the first neuropil, or to be transmitted to the second neuropil through higher-order neurons.

This difference raises interesting questions on the nature of this evolutionary change and its implications for the evolution of neural circuits and the homologies of photoreceptors and optic neuropils. One hypothesis is that the homologue of the long-projecting photoreceptors of *Parhyale* (most likely photoreceptor R5) has ceased to make long projections and terminates with the other photoreceptors in the first neuropil (Figure 8A), analogous to the changes observed in the ‘love spot’ of insects [70]. Alternatively, these photoreceptors may have been lost, indicating that all the photoreceptors of gammarid-like amphipods evolved from the short-projecting photoreceptors (R1-7) of their malacostracan ancestors (Figure 8B). These hypotheses would suggest that the first and second optic neuropils of *Parhyale* correspond to the lamina and the medulla respectively [31], separated by a chiasm as in other malacostracans and insects. The third discernible neuropil would be homologous with elements of the lobula complex. The lack of a chiasm connecting to this neuropil suggests it would correspond to the lobula plate [13].

**Figure 8.**
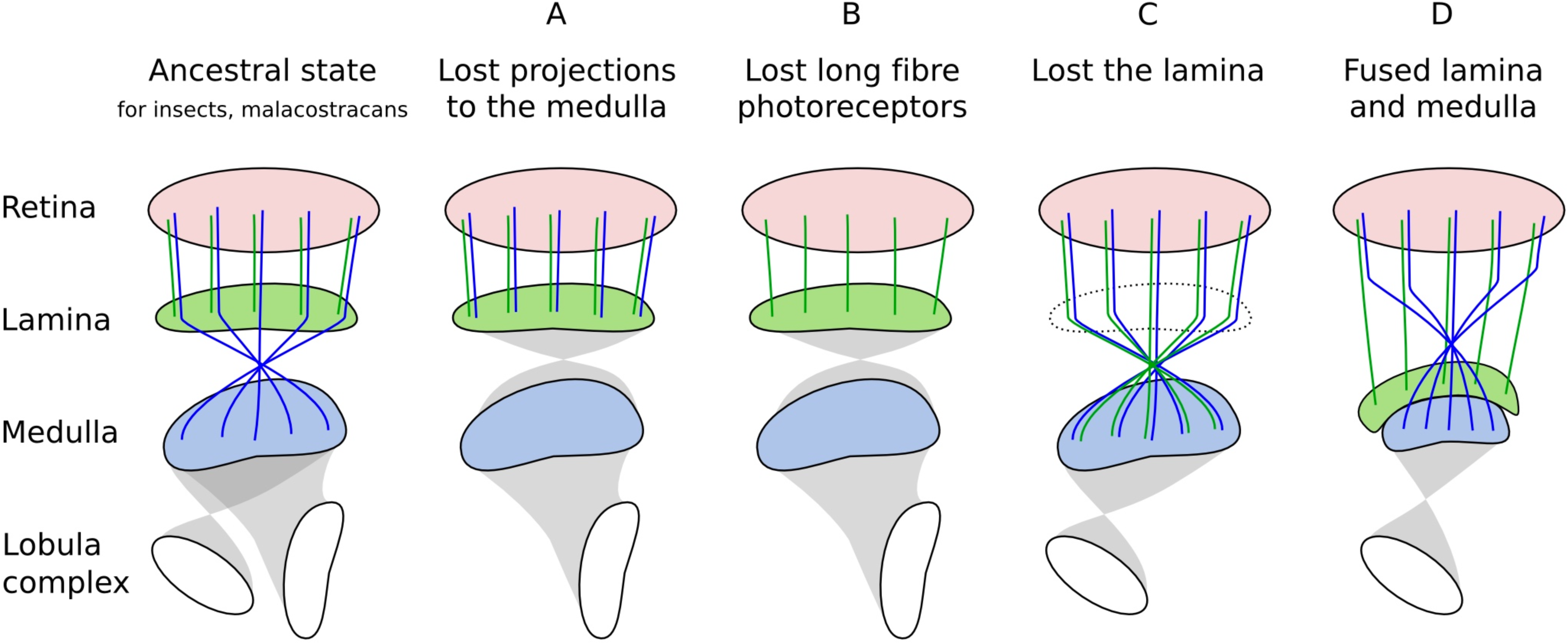
Hypotheses on the evolutionary changes that shaped the visual system of *Parhyale* Illustrations depict the ancestral state of insect and malacostracan visual systems and four alternative scenarios for the evolutionary origin and homologies of the photoreceptors and visual neuropils of *Parhyale* (explained in the main text). The suggested presence/absence of chiasmata in *Parhyale* is hypothetical. The retina, lamina and medulla are coloured red, green and blue, respectively; elements of the lobula complex (the lobula and lobula plate) are shown in white; short and long fibre photoreceptor types are coloured green and blue, respectively. Based on the presence/absence of a chiasm, we suggest that neuropil 3 of *Parhyale* is likely to be homologous to the lobula plate under hypotheses A and B. The second neuropil would correspond to the lobula under hypotheses C and D.

Altered photoreceptor projections could also reflect a more radical structural change, such as the loss of the lamina (Figure 8C) or the fusion of the lamina with the medulla (Figure 8D). This hypothesis would imply that the first optic neuropil of *Parhyale* corresponds either to the medulla or to the fused lamina and medulla, and the second neuropil corresponds to the lobula. In this case, the putative chiasm observed in the optic nerve of *Parhyale* would correspond to the chiasm preceding the medulla, and the chiasm between the first and the second neuropils would match the one found between the medulla and the lobula of insects and other malacostracans.

Whichever hypothesis is correct, it is tempting to speculate on whether these changes in optic lobe structure and photoreceptor projections could be associated with functional changes in the visual system of *Parhyale;* for example, with the release of functional constraints due to the dramatic reduction of spatial resolution, or the development of new light-driven responses based on polarised light. Detailed mapping of neural connections and behavioural assays for colour and polarisation sensitivity in different light regimes will be needed to resolve these questions.

Our work establishes *Parhyale* as a genetically tractable model for visual studies. In the near future, transgenic approaches will allow the genetic marking of single neurons, functional imaging using calcium sensors, and genetic manipulation of neural activity in this species, providing a foundation for exploring arthropod visual system development and function from an evolutionary angle.

## Materials and Methods

### *Parhyale* culture and handling

*Parhyale hawaiensis*, derived from a population collected at the Shedd Aquarium in Chicago [38], were kept in plastic boxes with artificial seawater (specific gravity 1.02) at 23-26°C. For embryo collection, adults were anaesthetised using clove oil (diluted 1:2500 in artificial seawater) for up to 20 min. Adults used for immunostaining or live imaging were anaesthetised in ice cold water for 30 min prior to fixation or imaging. Embryos were collected and staged as described previously [38, 71].

### Histological preparations

The heads of adult animals were fixed in 2.5% glutaraldehyde (Electron Microscopy Sciences #16300 or Agar Scientific R1020), 2% paraformaldehyde and 2% sucrose in Sorensen’ s phosphate buffer (pH 7.4) overnight at 4°C. After rinsing for 1h in phosphate buffer, samples were post-fixed in 1% OsO_4_ (Electron Microscopy Sciences #19150) in phosphate buffer for 2 hours at 4°C, followed by several washes in phosphate buffer. Samples were then dehydrated in a graded ethanol and acetone series, and embedded in epoxy resin (Electron Microscopy Sciences #14120). Semi-thin (2-5 micron) sections were cut on a Leica RM2265 microtome with an histology diamond knife, stained with toluidine blue and observed on a Leica DM6000 light microscope. For dark-adapted samples, animals were kept in darkness for 1 hour before being fixed in a dark room.

### Measurement of ommatidial acceptance angles

Estimates of the acceptance angle of each ommatidium were obtained from the ommatidial anatomy. In geometrical optics, the acceptance angle of a receptor is the angular width it subtends at the nodal point [3]. Because the crystalline cone is the single potentially focusing structure in the ommatidium, and the eye is in water with aqueous media on all sides, it is reasonable to assume that the nodal point is close to the centre of the crystalline cone. Determining the geometric position of the receptor is complicated because in reality the rhabdom is not a single plane but a volume shaped like a tapering cone. A single plane that would detect light within the same angle as the rhabdom would have to be between the distal and proximal ends of the rhabdom. Because the rhabdom is wider distally it is reasonable to assume that such a plane is closer to the distal than the proximal end of the rhabdom. If we assume that the plane with equivalent acceptance angle is located 1/3 of the distance down the rhabdom, then we can measure the angle that the rhabdom width at this position subtends at the assumed nodal point in the middle of the crystalline cone (Figure 1G). Performing such measurements on a number of good longitudinal sections of ommatidia provides acceptance angles in the range 15° to 30°.

### Transmission electron microscopy

Samples were fixed as described above, substituting phosphate buffer for 0.1 M sodium cacodylate buffer (Agar Scientific AGR1103) and embedded in epoxy resin (Agar Scientific AGR1031). Ultra-thin (50 nm) sections were cut using a Leica EM UC7 ultratome with a diamond knife. The sections were mounted on pioloform-coated copper grids, stained 30 minutes in 2% uranyl acetate and 4 minutes in Reynolds’ lead citrate [72], and examined at 100 kV using a JEOL JEM-1400 Plus transmission electron microscope. Micrographs were recorded with a JEOL Matataki CMOS camera using the TEM Centre for JEM1400 Plus software.

### Identification of opsins

The protein sequence of *Drosophila* Rh1 was used as a query to search the *Parhyale* embryonic and head transcriptomes by BLAST [37, 47, 48]. The best matches were confirmed to be opsins based on BLAST searches on the NCBI GenBank database and their predicted transmembrane structure (performed on PRALINE, [73]). The *PhOpsin1* and *PhOpsin2*. sequences thus identified were deposited in the GenBank/EMBL nucleotide sequence database.

The protein sequences of PhOpsin1 and PhOpsin2 were aligned with a set of 57 r-opsins from arthropods (and additional sequences from outgroups) which had been used previously to reconstruct the pancrustacean opsin phylogeny [49]. Most of the opsins in that dataset have known spectral sensitivities, helping to characterise the spectral properties of different opsin subfamilies. The protein sequences were aligned using the online tool MAFFT [74] using default parameters (scoring matrix BLOSUM62, gap opening penalty 1.53, strategy L-INS-i). The alignment was trimmed with TRIMAL [75] and AliView [76] to remove positions with multiple gaps (threshold value 0.11) and poorly aligned N- and C-terminal regions, respectively. The trimmed protein alignment contained 450 aligned residues. Based on this alignment, we reconstructed the opsin phylogeny using Maximum Likelihood implemented in IQ-TREE [77], using the best substitution model selected by the software. Branch support values were estimated from 1000 bootstrap replicates.

### In situ hybridization

Fragments of *PhOpsin1* and *PhOpsin2* genes were cloned from cDNA prepared from *Parhyale* embryos and adult heads. cDNA was prepared using oligo(dT) primers and the SuperScript III reverse transcriptase (Invitrogen) following the manufacturer’ s instructions. The fragments of *PhOpsin1* (1.4 kb) and *PhOpsin2* (0.6 kb) cDNAs were amplified by PCR using the following primer pairs: 5’ - GATTGGTTCTGCACGTGGC-3’ and 5’ -TTGAGTGACAACGTTTGTTGTCGG-3’ for *PhOpsin1* (primer annealing at 55°0, and 5’ -ATGTCCCACAGCCACAGCCCAT-3’ and 5’ -TCCGGAATGTAGCGGCCCCAGC-3’ for *PhOpsin2* (primer annealing at 62°C). The fragments were cloned into the pGEM-T Easy vector (Promega) and sequenced.

Antisense DIG-labelled RNA probes for *PhOpsin1* and *PhOpsin2* were made by *in vitro* transcription using the SP6 RNA polymerase (Promega) and the DIG RNA Labelling Mix (Roche), according to the manufacturer’ s instructions. As templates we used the *PhOpsin1* and *PhOpsin2* plasmid clones (described above) linearised by digestion with NcoI.

*In situ* hybridization was carried out in *Parhyale* embryos and adult heads following an established protocol [78]. Embryos were dissected from the eggshell and heat-fixed by submersion in boiling heat-fixation buffer (0.4% NaCl, 0.3% Triton X-100) for 2 seconds, immediately cooled by adding ice-cold buffer and standing on ice [79]. Adults were anesthetised and heat fixed following the same procedure. Brains were then dissected from the head capsule. All samples were then re-fixed in 4% formaldehyde in phosphate buffered saline (PBS) at 49C overnight. Hybridization was performed at 65°C, with a probe concentration of 3 ng/µl.

Stained specimens were washed in 100% methanol, propylene oxide (Electron Microscopy Sciences #20401) and embedded in epoxy resin (Electron Microscopy Sciences #14120) as described previously [80]. Specimens were then sectioned on a Leica RM2265 microtome with an histology diamond knife at a thickness of 4-5 microns and stained with eosin/erythrosin or basic fuchsin and observed on a Leica DM600 light microscope.

### Immunostainings

Antibody stainings were performed on embryos, vibratome sections of adult heads and dissected adult brains. Embryos were heat-fixed as described above. For vibratome sectioning, entire animals were fixed in Bouin’ s solution (Sigma) for 24 hours and washed several times in PBS with 1% Triton X-100 until the yellow colour disappeared. Fixed specimens were submerged in melted 3% agarose (in PBS). Once cooled, the block of agarose was trimmed, glued to the vibratome stage and placed in a PBS bath. 150 to 300 µm thick sections were made on a 7550 PSDS Integraslice vibrating microtome (Campden Instruments) and kept in PBS until staining. The sections were treated with 3% H_2_O_2_ and 0 5% w/w KOH in water [81] to remove eye pigmentation. For brain dissections, adult heads were fixed in 4% paraformaldehyde in PBS, overnight at 4°C. After several washes in PBS with 1% Triton X- 100, fixed heads were dissected from the ventral side and the brain was removed from the head capsule. All types of samples (embryos, sections or brains) were then washed in PBS with 1% Triton X- 100 (4 × 20 minutes) and incubated in blocking solution (1% BSA and 1% Triton X-100 in PBS) for 1 hour at room temperature. Samples were incubated for 4 to 5 days with primary antibody and 3 days in secondary antibody at 4°C, with four 30-minute washes in PBS with 0,1% Triton X-100 after each antibody. Nuclei were stained by incubating the samples with 0.1 mg/ml DAPI (Sigma). After an overnight incubation in 50% glycerol, samples were placed in Vectashield antifade mounting medium (Vector Laboratories) and mounted for microscopy, using clay as a spacer to avoid crushing of the samples under the coverslip.

Primary antibodies: mouse monoclonal 6-11B-1 for acetylated tubulin (diluted 1:1000; Sigma), mouse monoclonal 3C11 for synapsin (SYNORF1) (diluted 1:100; Developmental Studies Hybridoma Bank), chicken polyclonal for GFP (diluted 1:1000; Abeam abl3970), and rabbit polyclonal for turboRFP/mKate2 (diluted 1:500; Evrogen AB234). Secondary antibodies (all diluted 1:2000): goat anti-mouse Alexa 647 (Invitrogen), goat anti-rat Alexa 654 (Invitrogen), goat anti-chicken Alexa 488 (Abeam abl50173), goat anti-rabbit Alexa 555 (Life Technologies).

### Transgenic reporters

A genomic fragment containing c/s-regulatory regions of the *PhOpsin1* gene was amplified from *Parhyale* genomic DNA using primers 5’ - [TAAGCAGGATC]CAAGGAATACAGAATATCTCTGAGATTA-3’ and 5’ - [TAAGCACTCGAG]ATTACTCACTGTTCTCGAAGATTT-3’ (primer overhangs shown in brackets, primer annealing at 55°C). This fragment, which includes the 5’ UTR of *PhOpsin1*, was placed upstream of the start codon of the EGFP-CAAX coding sequence and cloned into the *Minos*. transposon vector [35] to generate donor plasmid *pMi(PhOpsin1:EGFP-CAAX)*. Cloning was performed using the MultiSite Gateway Pro kit (Invitrogen), after adapting the Tol2Kit plasmid library [82] for use with the *Minos* vector (details given in [83]).

A genomic fragment containing the c/s-regulatory regions of the *PhOpsin2* gene was amplified from *Parhyale* genomic DNA using primers 5’ - [ATCGATACGCGT]ACGGCGCGACGGAACATTCTGCATCTTAGCTTGTGC-3’ and 5’ - [CCTCTGCCCTCTC]CACTGCCCATCCTGAGCCTCTTCACCTTGAGG-3’ (primer overhangs shown in brackets, primer annealing at 63°C). This fragment, which includes the 5’ UTR, the start of the coding sequence, the first intron and part of the second exon of *PhOpsin2*, was placed upstream of the T2A peptide and the mKate-CAAX coding sequences, and cloned into the *Minos* transposon vector [35] to generate donor plasmid *pMi(3xP3-DsRed; PhOpsin2:mKate-CAAX)*. Cloning was performed using the Gibson Assembly Kit (NEB; details given in [83]). The T2A peptide was used to separate the N-terminus PhOpsin2 from mKate-CAAX and ensure that the two are expressed as separate polypeptides [84].

Transgenic lines were generated by microinjecting 100-150 ng/µl of each donor plasmid with 100 ng/µl Minos transposase mRNA in 1-cell stage *Parhyale* embryos, as described previously [35, 85]. Injected embryos were screened from stage 26 to the end of embryogenesis on a Leica MZ16F stereoscope. Embryos expressing the fluorescent marker in the eyes were used as founders to establish stable transgenic lines for each opsin reporter.

We noticed that old transgenic adults carrying the *PhOpsin1:EGFP-CAAX* construct showed a tendency to have damaged eyes and a variability in the number of neurons expressing EGFP, suggesting that the reporter may give stochastic expression in adults or that some photoreceptors in these animals may degenerate. For imaging we selected young adults that were least affected.

### Image acquisition and analysis

For live imaging, transgenic embryos were mounted in a drop of 1% low melting agarose in filtered artificial seawater on the surface of a 35 mm glass bottom dish (Ibidi µ-Dish) with the eye facing the glass surface.

Confocal images were obtained on a Zeiss LSM780 laser scanning confocal microscope. For live imaging we used a Zeiss C-Apochromat 40x NA 1.2 water immersion objective, while immunostained samples were imaged using Zeiss Plan-Apochromat 40x NA 1.3 oil, Plan-Apochromat 63x NA 1.4 oil and LD LCI Plan-Apochromat 25x NA0.8 objectives.

Image data were handled using Fiji [86]; the Enhance Local Contrast (CLAHE) plugin was used to enhance contrast. The confocal images shown in Figures 4B, 4C, 5A, 5C, 6A and 6D were deconvolved using the Fiji plugin DeconvolutionLab2 [87]; point spread function was calculated theoretically by the PSF generator plugin [88]. The 3D rendering shown in Figure 6D was obtained using the ImageJ 3D viewer plugin.

A confocal image stack of an adult brain, immunostained for acetylated tubulin, was used for the 3D reconstruction of the *Parhyale* optic lobes shown in Figure 5G. Neuropils were labelled manually using the TrakEM2 Fiji plugin [89].

### Light adaptation and phototaxis assays

To study adaptation to light intensity, live adults were immobilised in a petri dish using surgical glue (Dermabond), kept in artificial seawater and imaged using a Leica M205 stereoscope. After dark adaptation, by keeping the animals in darkness for >30 minutes, lights were turned on and an image of the eye was taken every 30 seconds. These experiments were repeated several times during the day and at night.

For phototaxis experiments, a two-choice tube maze (T-maze) was built using dark PVC tubes whose walls were entirely covered with a white felt fabric; the internal tube diameter was 9 cm and the length was 52 cm. The maze was placed in a tank with seawater, such that the bottom ∼3 cm of the maze were immersed. The extremities of the maze were covered with either black felt fabric or a light diffuser made of silk paper. Light intensity was measured using a light meter (Sinometer LX1010B, JZK) placed just behind the diffuser. To vary light intensity, the light source (LemonBest LED bulb, 650 lm, 6500K) was placed at different distances from the maze. For each experiment, groups of 10 animals (a mix of males and females) were placed in the centre of the maze. The centre was then covered to block light. The number of animals found at either extremity (or in the middle) of the maze was counted after 5 minutes. The experiments were performed at several times during the day (between 9am and 7pm); the light conditions at each extremity were randomised.

## Authors’ Contributions

APR and MA designed the experiments. APR performed the experiments, acquired and analysed the data. OG performed the electron microscopy. NL and IS performed the histology. DEN determined the optical properties of the eyes. APR, IS, DEN and MA interpreted the results. APR and MA wrote the manuscript. All the authors read, revised and approved the manuscript.

## Acknowledgements

We thank Ben Hunt and Ezio Rosato for access to their unpublished *Parhyale* head transcriptome, Nick Roberts and Mathias Wernet for discussions on polarisation vision and behavioural assays, Carsten Wolff and Pierre Godement for advice on vibratome sectioning, François Lapraz for guidance on reconstructing the opsin phylogeny, Caren Norden for Tol2Kit plasmids, Jana Fuhrmann for support on unpublished aspects of this project, and Nikos Konstantinides, Carsten Wolff for numerous discussions, and Mathilde Paris, Chiara Sinigaglia, Cagri Cevrim and Marco Grillo for comments on the manuscript.

